# SuperDCA for genome-wide epistasis analysis

**DOI:** 10.1101/182527

**Authors:** Santeri Puranen, Maiju Pesonen, Johan Pensar, Ying Ying Xu, John A. Lees, Stephen D. Bentley, Nicholas J. Croucher, Jukka Corander, Erik Aurell

**Author notes:** Corresponding authors: Erik Aurell, Jukka Corander. These authors contributed equally: Erik Aurell, Jukka Corander.

## Abstract

The potential for genome-wide modeling of epistasis has recently surfaced given the possibility of sequencing densely sampled populations and the emerging families of statistical interaction models. Direct coupling analysis (DCA) has earlier been shown to yield valuable predictions for single protein structures, and has recently been extended to genome-wide analysis of bacteria, identifying novel interactions in the co-evolution between resistance, virulence and core genome elements. However, earlier computational DCA methods have not been scalable to enable model fitting simultaneously to 10^4^-10^5^ polymorphisms, representing the amount of core genomic variation observed in analyses of many bacterial species. Here we introduce a novel inference method (SuperDCA) which employs a new scoring principle, efficient parallelization, optimization and filtering on phylogenetic information to achieve scalability for up to 10^5^ polymorphisms. Using two large population samples of *Streptococcus pneumoniae*, we demonstrate the ability of SuperDCA to make additional significant biological findings about this major human pathogen. We also show that our method can uncover signals of selection that are not detectable by genome-wide association analysis, even though our analysis does not require phenotypic measurements. SuperDCA thus holds considerable potential in building understanding about numerous organisms at a systems biological level.

**Author Summary:** Recent work has demonstrated the emerging potential in statistical genome-wide modeling to uncover co-selection and epistatic interactions between polymorphisms in bacterial chromosomes from densely sampled population data. Here we develop the Potts model based approach further into a fully mature computational method which can be applied to most existing bacterial population genomic data sets in a straightforward manner. Our advances are relying on more efficient parameter scoring, highly optimized and parallelized open source C++ code, which does not rely on the computation-intensive polymorphism subsampling approximations used earlier. By analyzing the two largest available population samples of *Streptococcus pneumoniae* (the pneumococcus), we highlight several biological discoveries related to the survival of the pneumococcus and co-evolution of penicillin-binding loci, which were not uncovered by the earlier analyses. Our method holds considerable potential for building understanding about numerous organisms at a systems biological level.

## Introduction

Direct Coupling Analysis (DCA) emerged less than a decade ago and has opened up a new direction of biological research by demonstrating that large population based protein sequence analysis can be leveraged to make accurate predictions about protein structure[1-7]. DCA has been successfully extended to predict secondary and tertiary RNA structure[8], synergistic effects on fitness of mutations in the *E. coli* lactamase TEM-1[9], the fitness landscapes of HIV proteins[10], prediction of mutation effects from sequence co-variation[11], and to genome-wide epistasis analysis for bacterial population genomics[12]. Our focus here is to significantly extend the applicability of DCA methodology by enabling scalable inference for two orders of magnitude larger than previously modeled dimensionality of sequence positions.

Maximum likelihood inference for the Potts models employed in DCA is intractable due to the form of the normalizing constant of the model distribution, hence various weaker criteria or approximations have been used to derive estimators of the model parameters. Notably, maximum pseudolikelihood is a statistically consistent inference method which has typically outperformed variational methods[13], such as the mean-field estimator[14]. The different software implementations based on regularized maximum pseudolikelihood for DCA applications (plmDCA)[3,14-17] have been designed for at most 1000-2000 sequence positions, after which the computation times tend to become prohibitive for practical purposes.

To enable use of plmDCA in a much higher dimensional setting, with the order of 10^5^ polymorphisms in a bacterial genome, Skwark et al.[12] stratified a genome into non-overlapping windows and sampled randomly a single SNP from each window to form haplotypes of approximately 1,500 sequence positions, on which the plmDCA implementation by Ekeberg et al.[15] could be directly applied. They then used a large number of repeated random sampling of positions from the stratified genome to aggregate information about interactions between polymorphisms across the genome. While this approach was demonstrated to successfully capture both known and novel interactions, it remains very computationally intensive and may still leave important interactions undiscovered as only a fraction of all possible combinations of interactions will be covered even when using large numbers of repeated samples. It is also a hybrid method which does not fully implement global model learning which is a conceptually central point of DCA. To avoid these problems, here we introduce a method termed SuperDCA, which can perform inference simultaneously for all SNP positions in a much higher dimension. These advances are based on a new computational architecture exploiting efficient parallelization and optimization to achieve scalability for up to 10^5^ polymorphisms. In addition to being significantly faster with more modest computational resources, we also show that the global inference with SuperDCA allows the discovery of previously undetected epistatic interactions that inform our understanding of bacterial biology related to survival of the pneumococcus at lower temperatures. SuperDCA is freely available from https://github.com/santeripuranen/SuperDCA

## Results

### Results of SuperDCA and comparison with genomeDCA

The Potts model for genome-wide epistasis analysis was fitted to two largest existing pneumococcal population data sets using the SuperDCA method; the Maela[12,18] and Massachusetts populations[19]. Two variants of the Maela population data were considered: one with only bi-allelic SNPs (81045 loci), filtered as in Skwark et al.[12] in order to maintain compatibility for comparison of the results, and the second with no restriction to bi-allelic SNP sites (94028 loci, **Methods**). For Massachusetts 78731 SNP loci were analyzed (**Methods**). Figure 1 shows the cumulative distributions of the estimated coupling strengths between SNP sites for the Maela and Massachusetts populations. In both cases a vast majority of the couplings were of negligible magnitude and could be discarded from further detailed investigation using the thresholds shown in Figure 1 (**Methods**).

**Figure 1:**
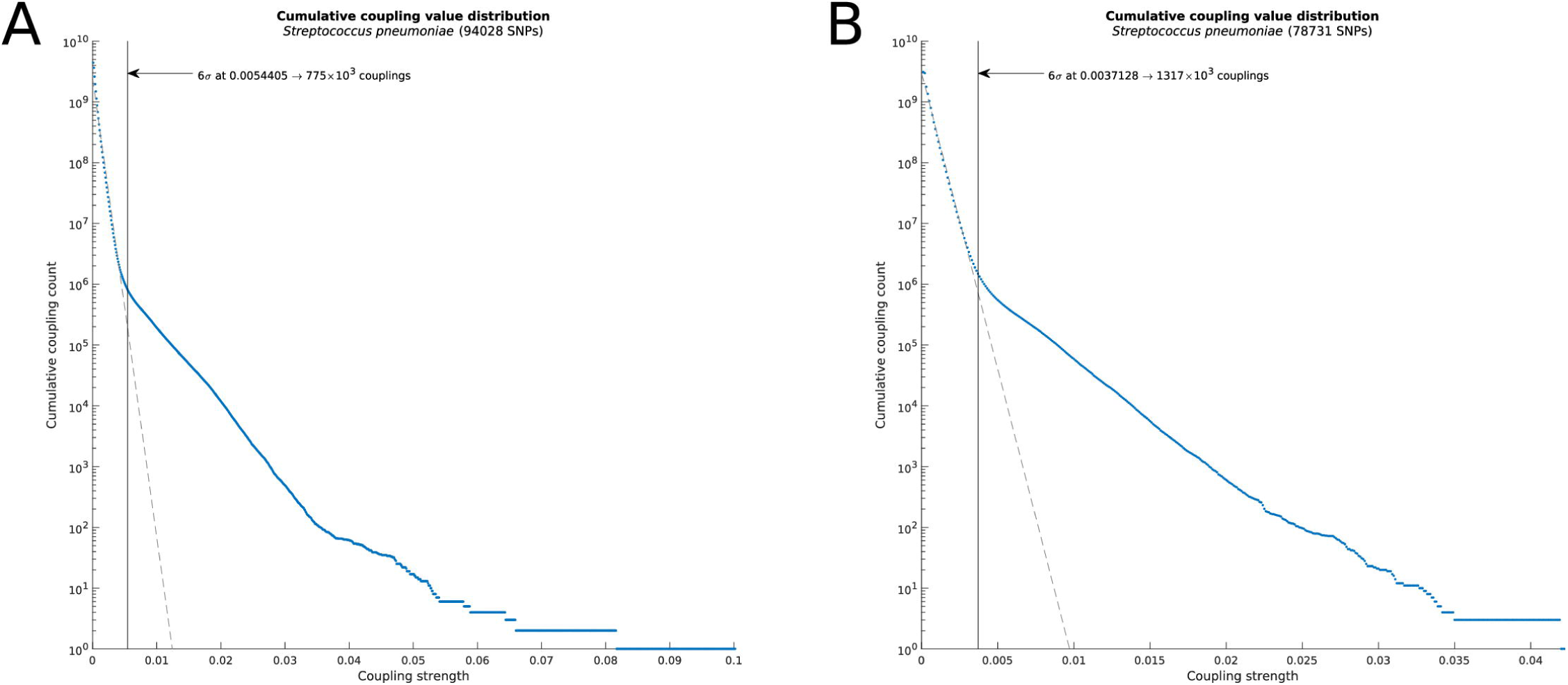
Log histograms of the cumulative distributions of estimated between-site couplings for Maela (left) and Massachusetts (right). The thresholds indicate the learned boundary between negligible and moderate to strong couplings.

Supplementary Figure 1 shows the overlap between the predicted genomeDCA and SuperDCA links on a gene level for Maela population. SuperDCA replicated the previously identified links between PBP gene pairs, as well as the network containing the *smc* gene. In contrast, SuperDCA did not identify significant links between pspA, divIVA, and the triplet upstream of ply, SPN23F19480-19500. In the simultaneous analysis which is not affected by chromosome stratification and random sampling of positions, the respective couplings no longer clearly deviated from the background dependence distribution, which is considerably wider for SuperDCA than for genomeDCA. This illustrated by a closer examination of the pairwise mutual information (MI) values (for further details see **Methods**) between the SNP loci in pspA, divIVA, and SPN23F19480-19500. The few stronger pairwise dependencies between the three genes disappear when all SNP loci are considered simultaneously. As a consequence of performing a full DCA analysis, in contrast to only partial DCA, the SuperDCA approach is less susceptible to highlighting weaker dependencies than genomeDCA.

### Epistasis in the penicillin-binding proteins

Since the bulk of the biological signal of between-site variation dependence presented in Figure 1 is due to linkage disequilibrium (LD) between sites in close proximity, we used a refined version of the phylogenetic ranking of the couplings (Supplementary Tables 1-3, **Methods**), to focus on the strongest candidates of co-selected loci. Figure 2 shows two sets of SNP loci which are involved in the top ranking couplings in Maela, alongside with the phylogenetic distribution of the alleles. The very top ranking couplings are between sites in the three penicillin-binding proteins (PBPs), as discovered in the earlier epistasis analysis which stratified the genome into non-overlapping windows and used the Potts model for sampled subsets of loci to reduce dimensionality[12].

**Figure 2.**
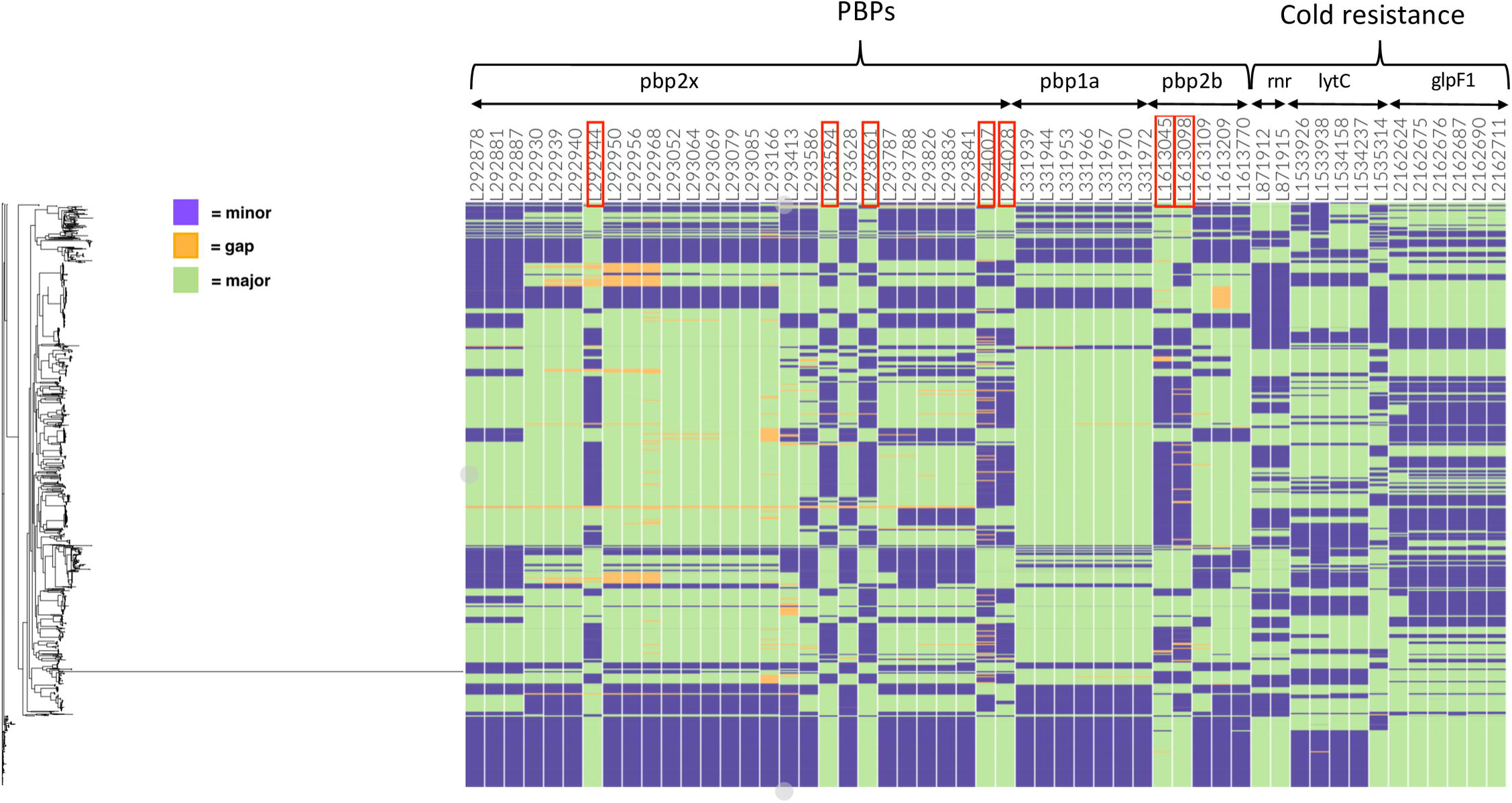
Maela population distribution of alleles at top ranked coupled SNP sites. The estimated genome-wide maximum likelihood phylogeny is shown on the left. Each column is labeled by the genome position, gene name and a corresponding functional categorization. Columns marked by red rectangles indicate coupled sites in *pbp2x, pbp2b* that have a reversed minor/major allele distribution compared with the remaining displayed SNPs in the same genes.

Figure 2 reveals a particular pattern of dependence between PBP mutations that adds significant biological information to the earlier findings[12]. The SNP positions marked by red rectangles in Figure 2 have an approximately reversed distribution of minor/major alleles in the population, which may reflect fitness differences regarding co-evolution of emerging mutations. In *pbp2x* the first marked position (codon position 359) corresponds to a synonymous mutation coding for amino acid phenylalanine, part of a conserved cluster of hydrophobic residues (Figures 3A and 3C) consisting of F353, P354, F393, L402, L403, and the E357 to K406 charge interaction located at the upper part of the transpeptidase domain near the active site. This cluster of residues likely has a role in maintaining structural integrity in this region (marked with cyan), as it is positioned next to the more mobile loop (marked with red) at residue positions 362-383 that partially covers the active site. Selection pressure seems to act in favor of the phenylalanine phenotype, since the genotype space clearly is explored here and switching the phenotype to the similarly sized and hydrophobic (but in contrast to phenylalanine non-aromatic) residues leucine or isoleucine, would only require a single non-synonymous mutation.

**Figure 3.**
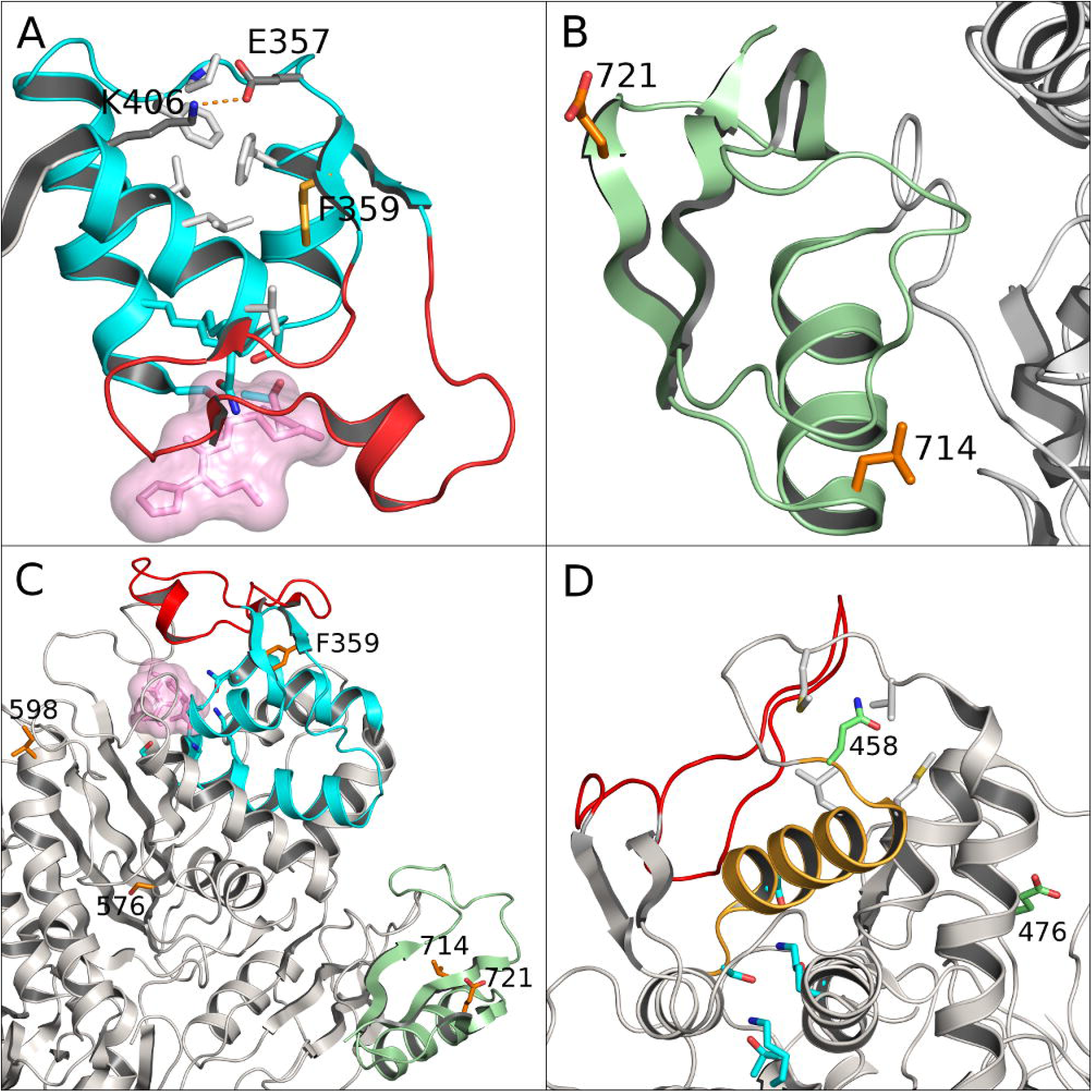
Structural mapping of the *pbp2x* (panels A-C) and *pbp2b* (panel D) positions marked in Figure 2. The panels show the transpeptidase domains of each PBP with active site residues shown in cyan and positions marked in Figure 2 as sticks in orange or green. Panel A depicts a structure-stabilizing cluster of conserved hydrophobic residues (light gray sticks) and charge interaction (dark gray) in a region proximal to (cyan cartoon) the *pbp2x* active site (with bound inhibitory antibiotic as pink space-filling volume) and a mobile loop (red cartoon) covering the active site. Panel B depicts the PASTA-2 domain essential for divisome complex function (green cartoon) with the bulk of the protein to the right (gray cartoon). Panel C shows an overview of the *pbp2x* transpeptidase domain colored as in the detail views in panels A and B. Panel D depicts the *pbp2b* transpeptidase domain region proximal to the active site with a helix (orange cartoon) mechanically connecting the active site to the ‘top’ of the protein. An adjacent mobile loop covering the active site is shown in red.

The second and third mutations (codon position 576, N/S/H amino acid changes; codon position 598, I/V amino acid changes) are conservative changes (Figure 3D) that may remotely affect the active site geometry or substrate association/dissociation kinetics, possibly as a compensatory mechanism for changes elsewhere. Active-site reshaping is an established cause of beta-lactam resistance in *S. pneumoniae*, where the involved polymorphisms can appear quite subtle at first sight. Our LD adjusted coupling scores indicate a very strong coupling between genome positions 294028/293661 in *pbp2x* and 1613045/1613098 in *pbp2b*. The fourth and fifth mutations (codon position 714, conserved L amino acid; codon position 721, E/Q amino acid change) are located in the PASTA-2 domain (Figure 3B; marked with green). The Q721 variant is prevalent in beta-lactam susceptible- and E721 in non-susceptible isolates. PASTA (PBP and Serine/Threonine kinase Associated) domains typically bind beta-lactams, however, a direct mechanistic role for 721 in beta-lactam resistance seems unlikely due to the structural position facing away from the protein core region. Rather, 721 is more likely to be involved in divisome complex formation and functions in a way that supports bacterial resilience in the presence of antibiotics; *pbp2x* and the PASTA domains therein are essential for bacterial division[20,21]. The characteristics and placement of L714 and the fact that all polymorphisms at this site are synonymous, point to a role in assuring structural integrity rather than in direct beta-lactam interaction.

In *pbp2b* the second marked position (codon position 458, D/N amino acid change) is located such that it may affect the active site in a mechanistic way via two distinct routes, either by indirectly modifying stability of the loop region (marked with red) proximal to the active site, or by slightly affecting the geometry of active site residues through the helix from 445 to 456 (marked with orange) directly connected to active site residues N445 and S443. The first marked position in *pbp2b* (codon position 476, G/E amino acid change) is spatially separated from 458. Although glycine at this site is more prevalent in beta-lactam non-susceptible- and glutamic acid in susceptible isolates, the potential role of the residue at this position in resistance remains unclear and would be a target for further experimental work.

Figure 4 shows a clear overlap between the Maela and Massachusetts populations in terms of identified links between genes involved in antibiotic resistance. For the two PBP gene pairs *pbp2x-pbp2b* and *pbp2x-pbp1a* the numbers of strong links between SNPs are large in both populations. For the pair *pbp1a-pbp2b* there is a pronounced asymmetry in this respect, such that the Massachusetts population harbors a large number of links whereas there are only very few in Maela. The latter observation is in line with the findings by Skwark et al.[12] which indicated that most interactions found between the PBP genes were between *pbp2x-pbp2b* and *pbp2x-pbp1a*. The fact that the Massachusetts population clearly deviates from this suggests that the co-evolution of PBPs may follow a non-congruent route in different populations. In the case of Massachusetts versus Maela, this may be a consequence of markedly different serotype distribution in the two populations, or other ecological constraints such as the varying selection pressure from different beta-lactam antibiotic usage. In Maela, beta-lactam prescriptions were almost exclusively amoxicillin, whereas in Massachusetts the pediatric prescription practice is likely to have been considerably more varied. Similar to the asymmetry of the extent of *pbp1a-pbp2b* couplings, the reverse allele distribution pattern discussed previously for Maela was not observed in the Massachusetts population. Given these differences our results suggest that the co-selective pressure on PBP gene polymorphisms acts differently depending on the type of the beta-lactams used in the population, warranting further experimental work to elucidate the mechanistic role of the coupled variations.

**Figure 4.**
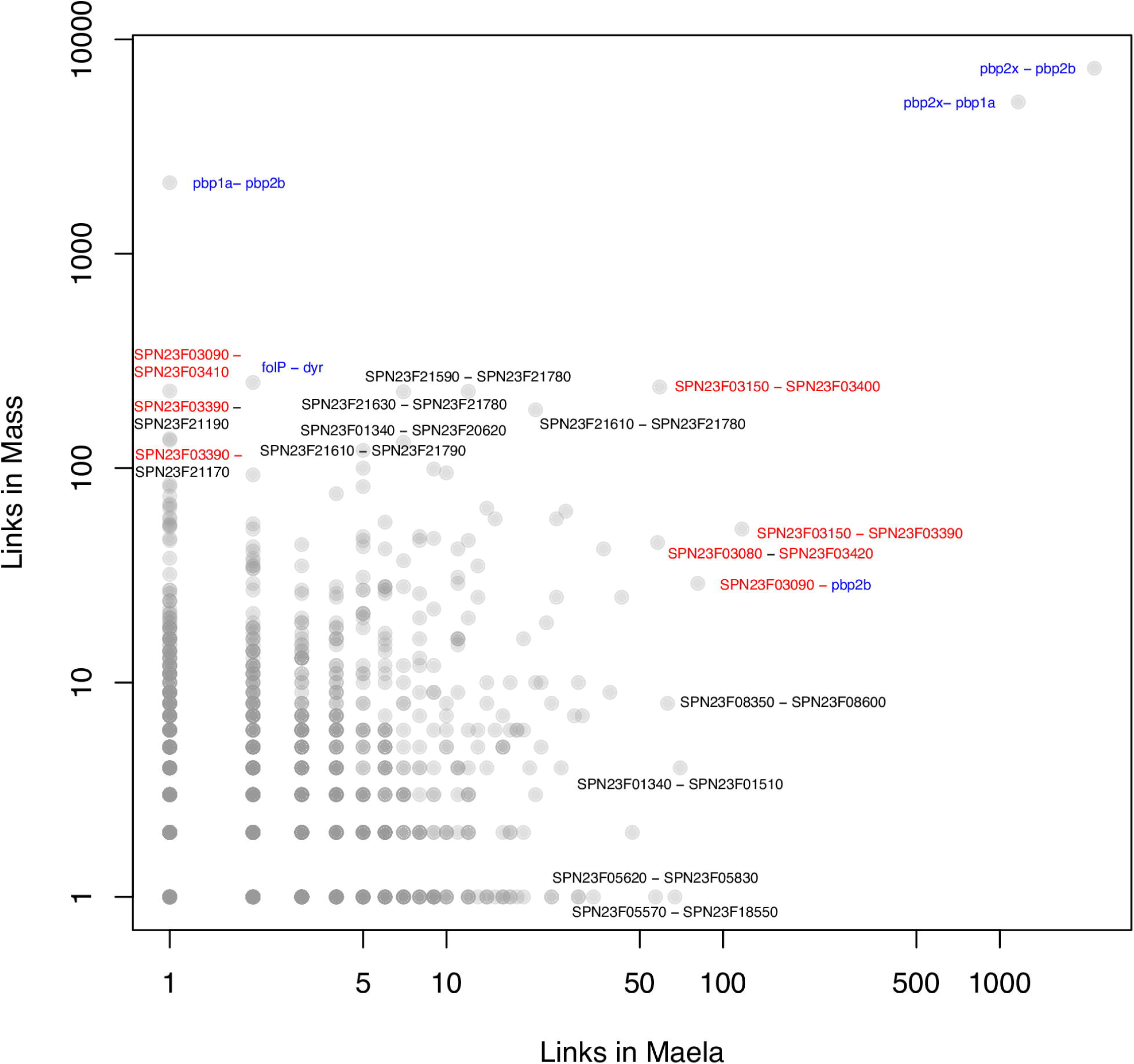
Overlap of estimated SNP interactions between the Maela and Massachusetts populations. Each dot represents an estimated link (interaction) between two coding sequences (CDSs), the blue CDSs are involved in antibiotic resistance, and the red CDSs are in close proximity to antibiotic resistance loci. Grey dots represent other functional categories not displayed here explicitly for visual clarity. Both axes are on log-scale and the values represent numbers of links in each CDS pair.

### Epistasis in cold tolerance and transmission potential

The current analysis additionally highlights several important between-site dependencies not identified by genomeDCA, showing greater sensitivity for identifying putative epistatic interactions. Firstly, the highest ranked SuperDCA couplings included twenty links between cold resistance-related genes exoribonuclease R (*rnr*), glyceroporin (*glpF1*), and lytic amidase C (*lytC*) (Figure 2), the strongest of which was ranked 668. In total, among the 5000 highest ranked couplings, there were two links between *glpF1* and *rnr,* and 18 links between *glpF1* and *lytC*. *GlpF1* is a transporter than imports glycerol, is involved in maintaining membrane fluidity with temperature changes[22]. The *glpF1* gene is at the 3’ end of its operon, with a tightly-folding BOX repeat at its distal end[23]. This would make the corresponding mRNA a potential target for *rnr*, a cold shock response 3’->5’ exonuclease that degrades tightly-folded RNAs that might be misfolded at lowered temperatures. Hence these interactions may be involved in tuning the expression of *glpF1* at lowered temperatures. Like *glpF1, lytC* is involved in maintaining the cell surface at lower temperatures, as it is the cellular amidase specialized at degrading peptidoglycan at lower temperatures (30 degrees Celsius, rather than 35-37 degrees Celsius)[24].

Previous work has demonstrated a significant seasonality in the transmission dynamics for the Maela population while carefully controlling for viral epidemics; the probability of the transmission being higher during the cold and dry winter months in comparison to warmer and more humid spring and summer months[25]. To examine whether the observed epistatic links related to survival at lower temperatures are connected with the seasonal transmission phenomenon, we examined the major and minor allele frequencies at the strongly linked cold resistance loci according to months, averaged over the three years 2007-2010 during which the data were sampled. Figure 5 shows clear temporal signals in terms of when the isolates carrying the linked minor/major alleles were sampled. The temporal changes in allele frequencies for the strongest cold resistance related link between *glpF1* (position 2162687) and *rnr* (position 871912), and also for the most strongly coupled sites between *lytC* (position 1533938) and *glpF1* (position 2162676), display a repetitive pattern of synchrony across years. In the first case, the proportion of major alleles in *glpF1* increases towards the end of the year, while in *rnr* the proportion of the minor alleles varies, being the dominant allele in January, April, and December. In the second case, the pattern in *glpF1* remains the same, but the proportion of minor alleles in *lytC* increases towards the winter months.

**Figure 5.**
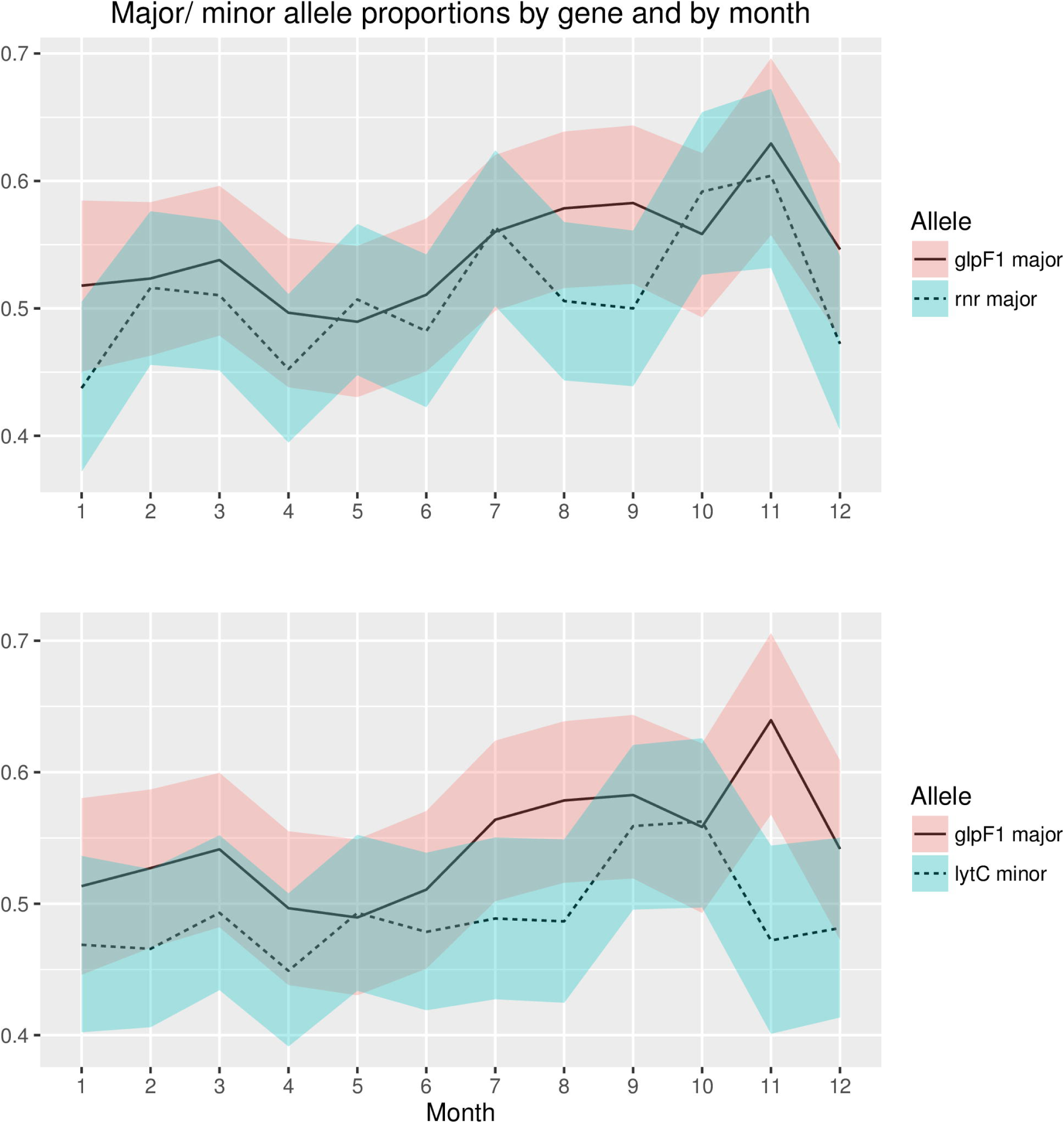
Seasonal variation of the allele frequencies for the two top cold resistance couplings between *glpF1-rnr* and *glpF1-lytC* averaged over three years, 2007-2010. The shaded areas indicate 95% confidence intervals.

These findings combined with the earlier results on Maela hosts being more susceptible for transmission during the cold and dry winter months[25], suggest that the recurrent selective advantage related to increased cold tolerance to facilitate survival outside hosts has been sufficient to shape the variation in population allele frequencies. To investigate whether the selection pressure on cold resistance genes could be discovered using a genome-wide association study (GWAS) approach, we coded the phenotype of each sample as winter or summer depending on the sampling date (**Methods**). We then applied the SEER GWAS method to identify polymorphisms that explain the variation in the phenotype[26]. Supplementary Figure 2 shows the Manhattan plot of the SEER analysis based on the annotated reference genome. No clear association signal can be seen and the SNP loci within the cold resistance genes are not associated with any markedly smaller p-values than the level of background variation of the association signal.

No cold resistance related couplings were found among the top 5000 couplings in the Massachusetts population, which may represent the less variable environmental conditions to which children are exposed, and the sampling of isolates only during winter, rather than year-round. In contrast, the Maela refugee camp conditions are such that the changes in selection exposure are more directly influential.

### Filtering on phylogenetic information

Inferred couplings from DCA typically have to be filtered to remove those that refer to trivial or non-informative dependencies. In the protein-structure applications very strong couplings are inferred among close neighbors along the peptide backbone, and are usually removed after model fitting by a simple distance based cut-off. A related issue is sampling bias, which for protein-structure applications has been handled by a reweighting applied to each sequence[1]. In bacterial sequence data produced from a sample taken from a small area over a limited period of time, a further issue is clonal inheritance; the meta-population is in a state of flux, and for a short window of time may not fully relax to the postulated Potts model of DCA. To compensate for this problem we used a refined version (**Methods**) of the phylogenetic re-ranking of the coupling estimates introduced in Skwark et al.[12] To visualize its effect, we consider mutual information to characterize the strength of pairwise dependencies between SNP loci. MI is a widely used information theoretical measure of dependence between discrete-valued variables, and it has been a popular tool as part of bioinformatics methods for DNA sequence analysis[27-29]. Here we use MI to characterize the strength of pairwise dependence between SNP loci as a function of their ranked estimated couplings alone and a ranking based jointly on couplings and phylogenetic criteria. Figure 6 shows the distribution of inferred MI values (**Methods**) for the two rankings in both Maela and Massachusetts population. The PBP-related couplings are nearly universally associated with higher MI values, indicating their tighter co-evolution despite of the negligible level of background LD between the three PBP segments. The distributions of large MI values have a clear shift towards a higher rank for both Maela and Massachusetts populations, which succinctly demonstrates the usefulness of using a phylogenetic ranking of coupling estimates to highlight co-selected sites above the background LD. A comparison of MI distributions for PBP-related SNPs for the two populations revealed that Maela displays stronger dependencies between the PBP mutations than Massachusetts (Supplementary Figure 3).

**Figure 6.**
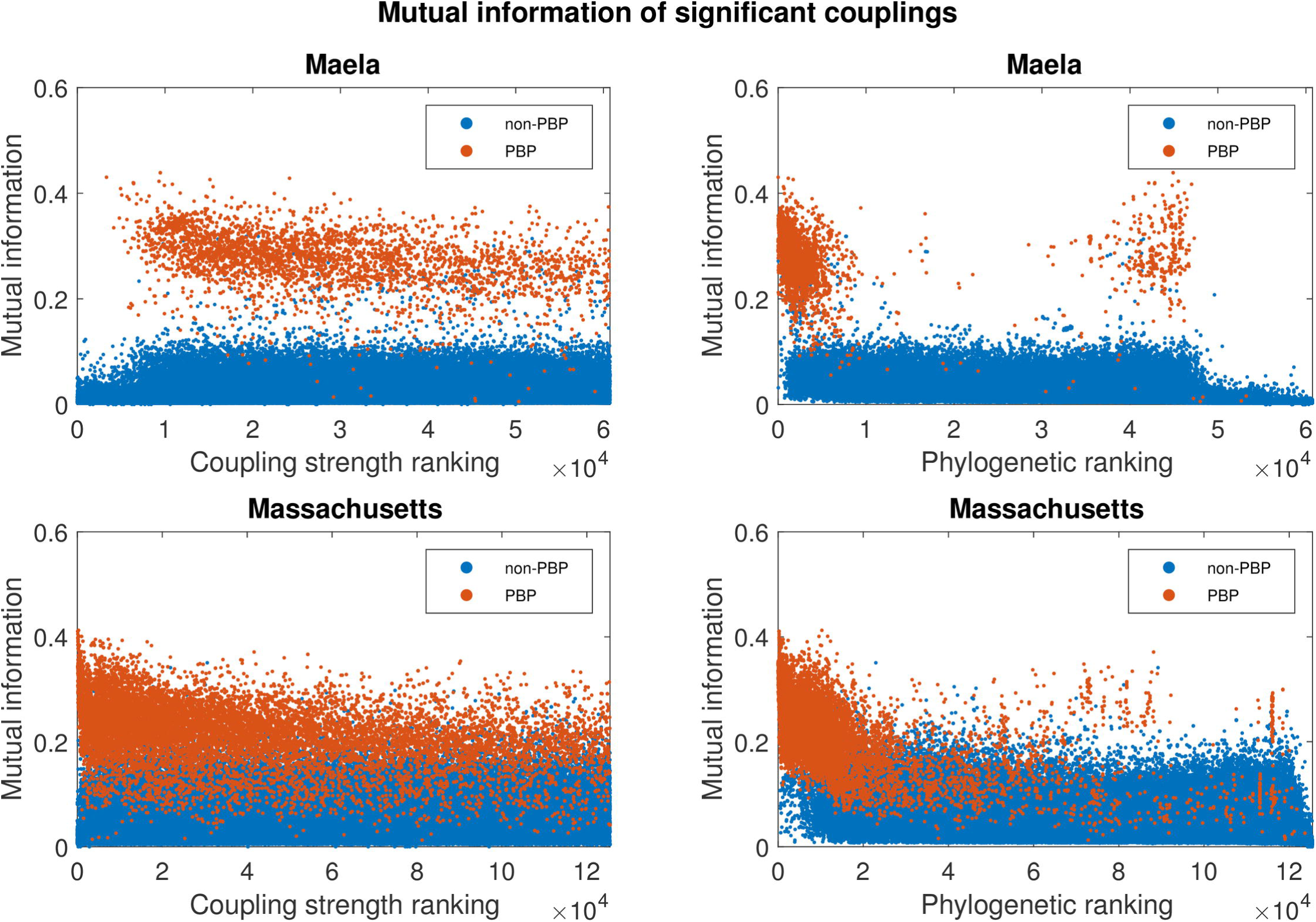
Estimated mutual information for 60749 pairs of SNPs (Maela) and 125469 pairs of SNPs (Massachusetts).

### Scalability improvements in SuperDCA

Overall, SuperDCA achieved an 18-fold effective performance increase over the earlier reference plmDCA implementation[15] on a single 20-core dual-socket compute node, enabling inference of 1.4*10^11^ parameters for a 94028 SNP genome dataset in less than 8 days, instead of an estimated 170 days. This was achieved through multiple alterations to the central algorithm explained below. Let (*S*_1_,*S*_2_…,*S_N_*) be a haplotype over *N* SNP loci, where each *s*_*i*_ can take values from an alphabet with cardinality *q*. Typically this cardinality varies between three (allelic states: minor/major/gap) and five (allelic states: A,C,G,T, gap). A Potts model assigns a probability distribution on such haplotypes defined by the following formula

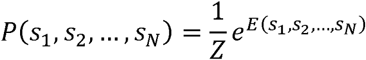

where the normalizing constant Z is known as the partition function and the expression in the exponent is

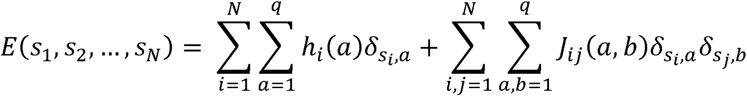

In above *δ*_*x.y*_ represents the Kronecker delta function which takes the value one if the arguments *x* and *y* are equal, and is otherwise zero. The linear terms are *hi(a) δ_si;a_* for different SNP loci and their alleles. The coefficients *h*_*i*_*(a)* parametrize a deviation from the uniform allele distribution for each SNP, independently of the values of all the other variables. The quadratic terms are the matrix elements *Jij(a,b) δ_si,a_δ_sj,b_* for different combinations of values of *i* and *j*, and *a* and *b*. The coefficients *Jij(a,b)* which are the *couplings* or *interactions* of pairs of SNPs, are defined as zero when the two indices i and j are equal. A coupling matrix with all elements equal to zero for non-identical locus index pairs implies that the alleles at these two loci are distributed independently in the population. Small positive values of the coupling matrix elements correspond to weak dependence between the SNP loci. In this paper we have addressed the issues of gauge invariance and gauge fixing in the Potts model[1] as described previously[12,15].

One of the major obstacles for using earlier plmDCA algorithms simultaneously on large numbers of SNPs without locus subset sampling is their large runtime memory requirements. plmDCA memory use is dominated by the storage of *q*^*2*^-dimensional parameter matrices *J*_*ij*_ where *q* is the cardinality of the SNP state space (the maximum value being *q* = 5 when a gap/indel is included). *J*_*ij*_ and *J*_*ji*_ are needed simultaneously for calculating the pairwise coupling value, and since the elements are inferred row- or column-wise for all *i* (or *j*) at a time, a straightforward implementation of the algorithm necessitates simultaneous storage of all couplings in an *N*-by-*N* matrix J, therefore storage is needed for *q*^2^(*N*^2^-*N*) scalar elements. The scoring of the estimated coupling matrices would then be calculated according to

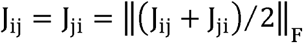

where *F* indicates the Frobenius norm. As an example, if a 10^5^ SNP genome alignment was characterized by 5-state alphabet and parameters stored in 64-bit floating point format, then the full interaction matrix J would require approximately 1.8 TB of memory, which is typically beyond the RAM available in state-of-the-art HPC cluster nodes. However, if the scoring of coupling values is instead calculated as

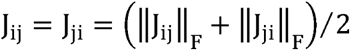

then runtime storage requirements are reduced by a substantial factor and the intermediate storage requirements for our example would shrink to 74 GB, which is well in the feasible range for current HPC nodes. Supplementary Figure 4 illustrates numerically that the above two scoring approaches lead to insignificant numerical differences in practice. SuperDCA uses this finding as one of its key improvements of plmDCA.

Performance profiling analysis identified high memory requirements and poor cache utilization as a major bottleneck for the performance in earlier plmDCA implementations when applied to higher-dimensional data. Parallel execution scaling also suffered due to memory bandwidth starvation. The maximization step was performed by Ekeberg et al.[15] using L-BFGS gradient-based optimization. However, the objective function required repeated traversal through all input data and the full parameter vector, emphasizing the need for an efficient data structure. To remedy these issues, a space-efficient, block-wise ordered data structure with simple state-pattern dictionary and run-length encoded indexing strategy for the genome data and a cache-friendly blocked memory layout for parameters were developed for SuperDCA and implemented in C++ (Supplementary Figure 5). A particular design choice was made to restrict the maximum value of *q* to 4. The resulting data structure reduced runtime memory use for nucleotide alignments by more than 4-fold compared with a typical dense data matrix representation. It also helped to reduce computing effort, improved processor cache utilization and enabled efficient utilization of SIMD vector instructions. The aggregate effect of these changes was an 8-fold improvement in single-threaded performance. The reduced main memory bandwidth use also helped improve node-level scaling as we measure a strong scaling factor of >0.7 up to 20 cores. Supplementary Figures 6-8 illustrate the computational scalability aspects for SuperDCA compared with genomeDCA.

## Discussion

Production of natural population genomic sequence data is currently still exponentially accelerating, highlighting the need for statistical methods that can generate detailed hypotheses for further experimental work regarding loci likely to be important in shaping bacterial evolution. Genome-wide association analysis has for a decade been the major general tool for such purposes, and more recently, its applicability to bacteria has been also demonstrated[26,30-32]. Skwark et al.[12] showed for the first time that statistical genome-wide modeling of joint SNP variation using DCA can uncover valuable information about co-evolutionary pressures on a large scale. This was done without relying on any phenotypic measurements, and using a hybrid scheme that does not fully employ the global model learning aspect of DCA. Here we built upon this initial observation to develop DCA into a powerful tool that is applicable to a majority of the existing bacterial population genome data sets in a computationally scalable manner. The biological insights on the differential evolution of PBPs, and the cold tolerance mechanisms, derived from the results of applying SuperDCA to two of the largest available pneumococcal genome data sets illustrate succinctly how such an approach could provide vital clues to the evolutionary processes under different ecological conditions in natural populations.

As the size of genome sequence data sets keeps growing, even our optimized parallel inference algorithm will eventually become too inefficient for practical purposes. Currently the chosen data and algorithmic architecture work extremely effectively for up to around 10^5^ polymorphisms. As bacterial whole genome alignments are typically of the order of 10^6^ sites, this should be sufficient for most population genomic studies. After this, the runtime will start to increase so rapidly that different computational strategies will be required for data sets including significantly more SNPs. Thus, an important topic for future research is to investigate how the Potts model inference can be performed in a reliable manner without resorting to a quadratic increase in the computational complexity as a function of the number of polymorphisms.

## Materials and Methods

### Data pre-processing

Bi- or tri-allelic loci with a minor allele frequency (MAF) greater than 1% were included in the analysis, provided that gap frequency was less than 15%. Gaps were not counted as alleles in the frequency calculations. To facilitate direct comparison with previous results[12] a separate dataset was prepared from the Maela input alignment using otherwise the same filtering rules, but for bi-allelic loci only. Filtering of 305245 SNPs in total resulted in two Maela input datasets for SuperDCA containing 94028 SNPs and 3042 samples using the former rules, and 81045 SNPs and 3145 samples using the latter rules. A subset of 103 samples containing mostly low quality reads were included in the data in the previous study, but were here removed from the source alignment prior to locus pre-selection for our 94028 SNP set. For the Massachusetts population the first set of filtering criteria resulted in 78733 SNPs and 670 samples.

### Hardware and inference details

Parameter inference was performed using a single 20-core HP SL230s G8 compute node with dual Xeon E5 2680 v2 CPUs and 256GB of DDR3-1667 RAM. Total wall clock run times were 186h (Maela with 94028 SNPs), 167h (Maela with 81045 SNPs) and 39h (Massachusetts with 78733 SNPs), including file I/O, pre-filtering and parameter inference. Weights correcting for the population structure, regularization and choice of hyper-parameters were calculated exactly as in the genomeDCA method[12]. Coupling estimates for the three data sets which exceeded the cut-off described below are provided as Supplementary Tables 1-3 at https://github.com/santeripuranen/SuperDCA.

### Prediction cut-off

The Potts models inferred in DCA are heavily over-parametrized. In protein contact applications the benchmark number of parameters is typically in the millions (number of residue pairs times q^2^, where q=21), while the number of samples varies typically from thousands to hundreds of thousands. Therefore only a small fraction of largest predictions are retained, commonly in the order of hundreds. Accordingly DCA manifests a varying degree of success when applied to protein families of the same size which has led to sustained efforts in algorithm optimization[6].

For the present and future applications to whole-genome data it is of more relevance to deliver a set of predictions at a pre-determined level of deviance from zero. An earlier approach using deviations from an extreme value theory distribution (Gumbel distributions)[12] was not applicable in the present setup, since we are not only sampling the tail of the coupling coefficients but estimate couplings for all possible pairs of SNPs. As shown in Figure 1, a semi-logarithmic cumulative distribution plot provides a computationally straightforward way to assess whether a particular coupling represents only random fluctuation near zero. The null distribution theory developed in Xu et al. for DCA inference procedures provides a strong motivation for using the linear part of the distribution near the origin as representation of the noise level signals[33]. To obtain a threshold we first perform a systematic scan over the histogram bins to fit a two-component linear spline function to the cumulative distribution. The standard deviation of the null couplings was then estimated using the part of the distribution between zero and the breakpoint. Similar to the Gumbel fit deviance level used by Skwark et al. [12] we then exclude all couplings that are less than 6 standard deviations away from the linear trend from further analysis. Figure 1 illustrates that this procedure effectively filters out the vast majority of all possible couplings as noise, and allows the downstream analysis to focus on the relevant signals.

### Phylogenetic ranking of estimated couplings

By default, SuperDCA includes gaps as a state in the Potts model if they are found in the alignment at sites fulfilling the SNP pre-filtering criteria. Some gaps can be considered informative, representing indels, while some simply relate to sites that are difficult to sequence. Hence some strong gap-induced couplings can represent lower quality sequence data instead of true between-site interactions, and they should be automatically de-emphasized to better enable assessment of the biological meaning of the inferred couplings. The superDCA coupling estimates are by default re-ranked using a combination of the three criteria described below, in addition to the actual value of the coupling.

Let C be a set of estimated couplings and 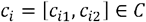 a pair of SNP loci represented by their genome position indices. Let 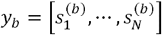 be a haplotype over the N SNP loci. Further, *S*_*i*,1_, is set of haplotypes carrying a minor allele at locus *C*_*i*,1_ and *S*_*i*,2_ a set of haplotypes with a minor allele at locus *C*_*i*,2_ The first phylogenetic ranking criterion is the minimum of the average genome-wide Hamming distances of all pairs of isolates 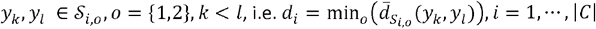 where 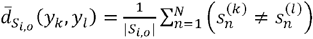.

Our second criterion is the normalized number of hierBAPS[34] clusters including isolates carrying the minor alleles at the two coupled loci, i.e. 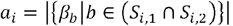, where *β*_*b*_ is the designated hierBAPS cluster for haplotype b. Finally, the third criterion is the percentage of isolates where both SNP loci involved in a coupling had the minor allele, i.e. 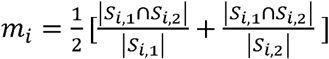

The above three criteria are normalized by: 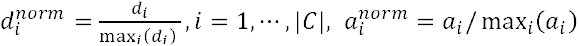 and 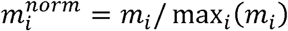,after which they are combined to a single ranking criterion 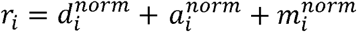 having a maximum value of three and a minimum equal to zero. Large values emphasize cases where both minor alleles at coupled loci are simultaneously widely distributed across the population. In cases where gaps at any two loci are phylogenetically spread in the population and would have led to a large estimated coupling values, they are still de-emphasized since they are not counted as minor alleles. The above criteria are derived by normalizing the individual coupling re-ranking measures developed by Skwark et al. [12]. The hierBAPS clusterings were obtained from the original publications introducing genome sequences for the Maela and Massachusetts populations[18,19].

### Mutual information calculations

Mutual information is an information theoretic measure of the mutual dependence between two variables. Let *X*_1_ and *X*_2_ be two discrete variables with outcome spaces indexed by *i*=1 … r_1_ and *j*=1, … r_2,_respectively (the outcome indexing differs here from the earlier description of Potts model for notational simplicity). Let *p=(p_ij_)* represent the joint distribution over the variables such that p_ij_ corresponds to the probability of (*X*_*1*_=*i,X*_*2*_=*j*). The mutual information between *X*_1_ and *X*_2_ is then calculated by

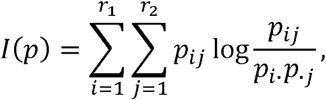

where 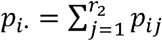 and 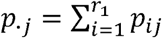 are the marginal probabilities for the corresponding variables (here SNPs). In practical applications the joint distribution is usually not known and must be estimated from data. The standard approach of estimating the probabilities is to use the maximum likelihood estimates given by the relative frequencies 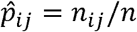 where *n*_*ij*_ denotes the count of the corresponding configuration and *n* is the sample size. A drawback of the standard frequentist approach is that it does not account for the uncertainty of the estimates.

In this work we adopted a Bayesian approach[35], where we put a prior density function *f(p)* on the *r*_*1*_.*r*_*2*_ unknown joint probabilities. Taking the data into account, the posterior density function *f*(p|data) can be calculated from the prior using Bayes’ theorem. The posterior density over the mutual information given some dataset is then obtained through

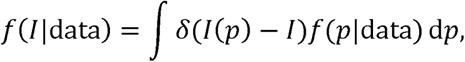

where *δ*(.)is the Dirac delta function.

The above density function can be approximated using a Monte Carlo simulation by sampling from *f*(p|data) In particular, assuming a Dirichlet prior on *p* with hyperparameters *α*_*ij*_ enables a straightforward sampling scheme since the posterior *p*|data then follows a Dirichlet distribution with updated hyperparameters 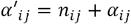. Our main interest is to calculate the Bayesian point estimate given by the posterior mean

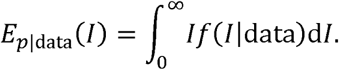

For this particular purpose, there exists an exact closed-form expression[35]:

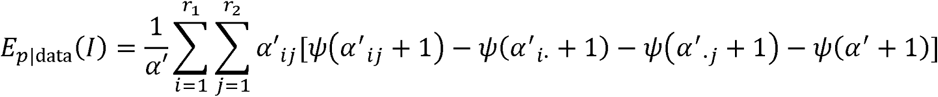

where Ψ(.) is the digamma function and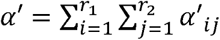.When using the above estimator we define the hyperparameters using the reference prior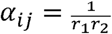. To adjust for the population structure in the sample, we use the same re-weighting scheme as was applied in our SuperDCA inference with a similarity threshold of 0.90. Finally, to remove the influence of gap-gap interactions, we did not include sequences for which either of the two considered loci had a gap value.

### GWAS for the seasonality phenotype

We coded season as a binary variable based on whether isolates were acquired during the winter or the summer. We then tested 123791 SNPs passing simple frequency filtering (>1% MAF) for association with this variable using SEER[26], which performs a logistic regression at every SNP. We used the first three multi-dimensional scaling components of the pair-wise distance matrix as fixed effects to control for population structure[26].

### Structural analyses

Crystal structures of *S*.*pneumoniae* PBPs with the following IDs: 2WAF (*pbp2b*), 1QMF and 1RP5 (pbp2x) were retrieved from the Protein Data Bank[36] (www.rcsb.org; accession date January 8, 2016) and visualized in The PyMOL Molecular Graphics System, Version 1.8.4.0 (Schrödinger, LLC). A chimera of 1QMF (chain A residues 257-618) and 1RP5 (chain A residues 64-256 and 619-750; missing sidechain atoms of E721 were reconstructed) was used for visualizing pbp2x.

## Supporting information legends

**S1 Fig. Overlap of the predicted genomeDCA and SuperDCA couplings in Maela population.** Lines are plotted between genes having at least three SNPs linked from the same genes, both in genomeDCA and SuperDCA. This results in 274 overlapping interactions. The thickness of lines is proportional to the number of linked positions within the corresponding genes. Gene annotations shown outside the circle are centered at the positions of the corresponding genes. Red labels are given for genes linked tightly both with genomeDCA and SuperDCA, black labels for genes linked only with genomeDCA and blue labels for genes linked only with SuperDCA.

**S2 Fig. SEER GWAS Manhattan plot of p-values for the winter/summer phenotype.**

**S3 Fig. Boxplots of MI value distributions for pairs of SNPs in two different PBP genes.**

**S4 Fig. Comparison of the norm-of-mean (vertical axis) versus the mean-of-norms (horizontal axis) summary score strategies.** Differences between the two are negligible for stronger, statistically significant coupling values and show more pronounced deviations only towards the sub-significant domain. The plot was calculated using a 25% uniformly random sample of loci from the 94028 SNP Maela dataset using the full data as background. Coloring marks coupling value count in log-scale.

**S5 Fig. Schematic drawings of the central data structures used in SuperDCA for storing input state data (nucleotide alignments) and the inferred parameters**. The *input data matrix* is stored such that samples (i.e. isolate genomes in our case) are ordered row-wise, with each sample divided into blocks of size *b*. Only column-wise unique blocks are stored. Block indices are stored for sample-oriented access to the data and sample indices for column- oriented access. Block index lists can optionally be run-length encoded, which leads to very significant space savings in particular when storing full-genome alignments with large regions of low column-wise variation. Index-lists for column- oriented access can similarly be collapsed for saving storage space when indices form contiguous (ascending) sequences. The *inferred parameters* are stored in blocked format such that all parameters relating to a particular column block in the input data are grouped together.

**S6 Fig. Comparison of SuperDCA versus plmDCA parallel scaling efficiency.** SuperDCA (blue curve) shows markedly stronger scaling than plmDCA (red curve). The scaling numbers were obtained as a mean of three runs of three 2-permil uniformly random samples of loci (188 loci) from the 94028 SNP Maela dataset and using the full data as background. Inferred parameter storage was disabled in plmDCA for the purpose of benchmarking. All benchmarks were run on a single 20-core HP SL230s G8 compute node with dual Xeon E5 2680 v2 CPUs and 256GB of DDR3-1667 RAM.

**S7 Fig. Comparison of SuperDCA versus plmDCA sample size scaling.** The sample-compressing datastructure used in SuperDCA enables markedly stronger scaling (blue bars) with increasing sample size than plmDCA (red curve). The scaling numbers were obtained as a mean of 9 runs: three-by-three sets of runs using a uniformly random sample of sequences and run for three 2-permil uniformly random samples of loci (188 loci) from the 94028 SNP Maela dataset and using the full data as background. See caption of Supplementary Figure 6 for details of benchmark hardware.

**S8 Fig. SuperDCA runtime improvement over plmDCA.** The single-threaded performance of SuperDCA is more than 8-fold that of plmDCA. Due to the greater parallel scalability of SuperDCA the performance delta grows as more compute threads are used, reaching more than 17-fold when run on 20 cores. See caption of Supplementary Figure 6 for details of benchmark settings and hardware.

